# Synergy between Enterobacteriaceae and dietary components shapes the competition between the dominant anaerobes of the human gut

**DOI:** 10.1101/2025.09.03.673633

**Authors:** Caroline Tawk, Youssef El Mouali, Kun D. Huang, Achim Gronow, Lisa Osbelt, Johanna Rapp, Nicola Segata, Hannes Link, Athanasios Typas, Till Strowig

**Affiliations:** Department of Microbial Immune Regulation, Helmholtz Centre for Infection Research; Braunschweig, 38124 Germany; German Center for Infection Research (DZIF), partner site Hannover-Braunschweig, Braunschweig, Germany; Interfaculty Institute for Microbiology and Infection Medicine Tübingen, University of Tübingen; Tübingen, 72076 Germany; Cluster of Excellence “Controlling Microbes to Fight Infections”, University of Tübingen, 72076 Tübingen, Germany; M3 Research Center, University of Tübingen, Otfried-Müller-Str. 37, 72076 Tübingen, Germany; Department of CIBIO, University of Trento, Trento, Italy; IEO, Istituto Europeo di Oncologia IRCSS, Milan, Italy; European Molecular Biology Laboratory, Molecular Systems Biology Unit; Heidelberg, 69117 Germany; Cluster of Excellence RESIST (EXC 2155), Hannover Medical School, Hannover, Germany; Center for Individualized Infection Medicine (CiiM), a joint venture of Hannover Medical School and HZI, Hannover, Germany

## Abstract

The human gut microbiota is frequently dominated by the Bacteroidota phylum, with Bacteroidaceae and Prevotellaceae families prevailing in modern and traditional lifestyles, respectively. While these distinct profiles have been well-documented across cohorts, the underlying mechanisms remain poorly understood. Here, we combine defined gut community models and genetic tools to experimentally dissect the ecological principles that favor Prevotellaceae or Bacteroidaceae. Testing 94 dietary components in the community model shows these two families compete for overlapping nutritional niches. Notably, *Segatella copri*, a key Prevotellaceae representative, outcompetes Bacteroidaceae strains when preferred polysaccharides are available. Interestingly, this competitive advantage depends on the presence of Enterobacteriaceae, which synergize with dietary components to promote *Segatella* dominance. Accordingly, we observe that global *Segatella*-rich non-Westernized microbiomes correlate with higher Enterobacteriaceae content. Our results suggest a revised hypothesis in which the ecosystem-level dynamic interplay between dietary components and key commensals within the community contributes to distinct Bacteroidota family dominance.

## Introduction

Profiling the gut microbiota of humans globally revealed diverse compositions of a healthy microbiota. ^1,2^. While the precise taxonomic composition varies significantly among individuals, distinct signatures for a healthy gut microbiota are now well recognized ^3–5^. These signatures have been described as enterotypes or bacterial guilds and are attributed to distinct lifestyles and dietary patterns ^4–6^. However, the factors that determine common microbiota signatures remain experimentally uncharacterized. Establishing clear causal links between intrinsic or extrinsic factors and microbiota signatures has proven challenging due to the inherent complexity and variability of the microbiome ^7^. The use of synthetic defined gut communities has emerged as a valuable approach for investigating microbial ecological interactions, such as cross-feeding and antagonism, as well as community capacities, including responses to drug perturbations, colonization resistance, and community resilience ^8–11^. Therefore, characterizing the components that influence the structure and stability of the microbiota is essential for understanding how variations in the microbiome contribute to disease.

Two widely observed gut microbiota signatures, shaped by the abundant Bacteroidota phylum, notably diverge at the level of bacterial families. Some individuals are dominated by Bacteroidaceae species, while others are dominated by Prevotellaceae species ^12^. Bacteroidaceae are prevalent mainly in Westernized populations, correlate with diets rich in animal fat and industrialization, and have been the subject of extensive investigation ^13,14^. Prevotellaceae, specifically members of the *Segatella copri* species complex (formerly *Prevotella copri*), are prevalent in non-Westernized populations, correlate with plant-rich diets and rural lifestyles, and are much less understood ^15^. Interestingly, dietary patterns alone do not fully account for the distinct microbiota signatures associated with these bacterial families, suggesting there are additional unidentified factors that influence their establishment ^13,16–18^. Notably, there appears to be an inverse correlation between the dominance of these two bacterial families, suggesting potential competition for similar ecological niches, likely driven by their shared preference for dietary glycans; however, this has not been thoroughly investigated experimentally ^19^. Bacteria belonging to these two families encode for a large repertoire of glycan utilization loci named polysaccharide utilization loci (PULs) and numerous Carbohydrate-Active enZYmes (CAZymes). *Segatella sp.* encode for an average of 25 polysaccharide utilization loci (PULs) and 40 Carbohydrate-Active enZYmes (CAZymes) predicted to mainly degrade plant-derived polysaccharides ^20,21^, whereas species in the Bacteroidaceae family often encode for hundreds of PULs and CAZymes specialized in the degradation of plant– and animal-derived polysaccharides^22^. Having such a large portion of the genome dedicated to diverse polysaccharide utilization suggests that complex carbohydrates may drive competition and community dominance in *Segatella sp.* and Bacteroidaceae species. However, the exact nature of the interactions and relationships between these prevalent gut bacteria remains unclear.

Here, we designed synthetic gut communities to resolve the specific interactions between *Segatella copri* and Bacteroidaceae species within a complex microbial network. We analyze the direct effect of 94 dietary compounds, including dietary glycans and vitamins, on these assemblies. We demonstrate that Prevotellaceae and Bacteroidaceae compete within the community, where *Segatella sp.* outcompete Bacteroidaceae depending on preferred dietary glycans and the presence of other commensals in the community. Specifically, we show that Enterobacteriaceae species largely promote *Segatella copri* within the community and cross-feed on monosaccharides from specific fibers degradation. In our metagenomic analysis of human cohorts, we found that Enterobacteriaceae are both prevalent and diverse in *Segatella*-rich, non-Westernized human microbiomes and foods. Together, our study reveals that the interplay between dietary fiber and Enterobacteriaceae shapes the competitive dynamics of dominant Bacteroidota in the gut.

## Results

### Supplementation with a single dietary component induces major shifts in a gut community model

Diets rich in plant-derived fibers have been shown to correlate with a high prevalence of Prevotellaceae in the microbiota ^18,21^. To determine the specific factors that drive the predominance of Prevotellaceae within a bacterial gut community, we designed a synthetic commensal community comprising 21 human gut isolates representing the four major phyla of the healthy human microbiota (Fig. 1a, Supplementary Table 1; see Methods). We tested a total of 94 dietary components, including 79 complex dietary glycans from algal, animal, bacterial, fungal, plant, and synthetic sources, as well as 15 essential vitamins for their effect on gut community structure and composition (Fig. 1b). We included vitamins in our analysis because of their important role as essential food components and their potential influence on bacterial competition within the gut community ^23^. The community was grown in the modified Gifu anaerobic medium (mGAM), a rich medium where the main defined carbohydrate sources are starch (0.5%) and glucose (0.05%). The glycans were added to mGAM at a concentration range of 0.04-0.2%, depending on their physicochemical properties, and the vitamins were added at their specific physiological ranges (Supplementary Table 2; See Methods). The communities were first assembled in mGAM, and the relative abundance of each member was quantified at the initial time point (Input), and at passage 3 (P3) once the community had stabilized, using 16S rRNA amplicon sequencing. Bacteroidaceae (represented by five species from the Bacteroides genus: *Bacteroides thetaiotaomicron, Bacteroides fragilis, Bacteroides uniformis, Bacteroides ovatus,* and *Bacteroides caccae*, and one species from the Phocaeicola genus: *Phocaeicola vulgatus*) predominated over Prevotellaceae (represented by *Segatella copri*) within the community in mGAM alone (Fig. 1c). Interestingly, supplementing the rich media with a single glycan or vitamin was sufficient to induce large changes in community composition (Fig. 1d). Most components did not significantly affect the final optical density or total cell number of the community after 24 hours of growth, except for vitamin C, which exhibited antimicrobial activity, and some gelling compounds, which affected the final OD600 absorption (Extended Data Fig. 1a,b). A majority of glycans (56/94), particularly from plant, algal, and bacterial origins, lead to a significant expansion of *S. copri* (Fig. 1d,e). Interestingly, for most of these compounds, *S. copri* expanded at the expense of Bacteroidaceae strains, shifting the balance toward a higher relative abundance of *S. copri* compared to the Bacteroidaceae strains (Fig. 1d). The addition of B vitamins (B3 or B5) was also sufficient to increase *S. copri* relative abundance in the gut community (Fig. 1d,e). Notably, the bacterial species whose relative abundance was significantly affected by most compounds belonged to the Bacteroidales order: *S. copri* and the Bacteroidaceae species, as well as *Enterocloster boltae* (Fig. 1f). Interestingly, the relative abundance of *S. copri* was increased by the largest number of compounds (n=56) (Fig. 1f). In contrast, the relative abundance of *P. vulgatus* was decreased by the largest number of compounds (n=63), followed by *B. ovatus* (n=46) and *B. uniformis* (n=42) (Fig. 1f).

**Figure 1.**
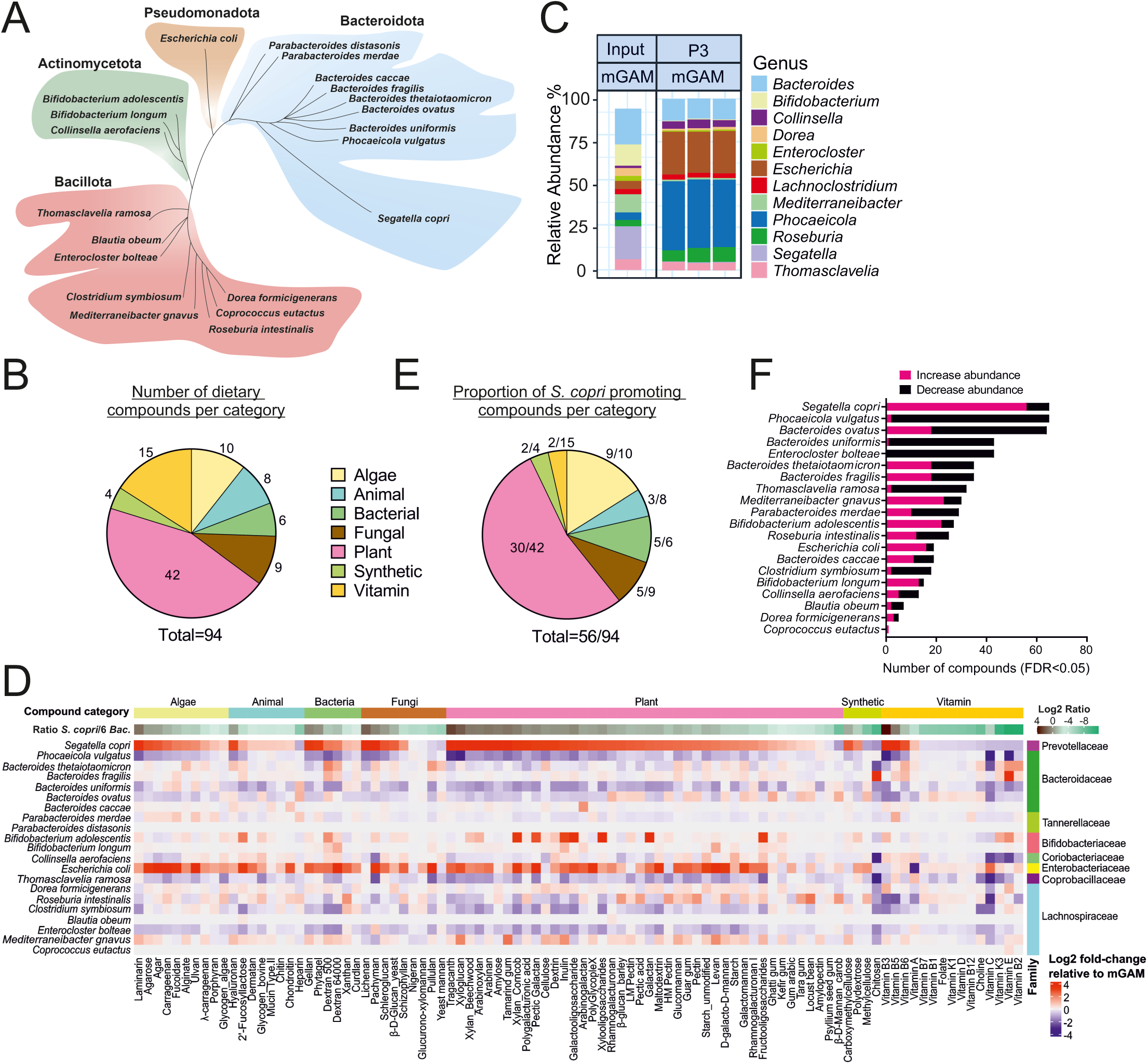
Food-derived complex carbohydrates and vitamins promote *Segatella copri* in a gut community model. (**A**) Phylogenetic representation of the 21 commensal human isolates comprising the defined gut community. (**B**) Pie chart representing the number of dietary compounds tested in each category. (**C**) Taxonomic bar plots showing the relative abundance of the top 12 community genera at passage 3 and the input in mGAM. Data is from 16S rRNA amplicon sequencing with three biological replicates shown. (**D**) Heatmap showing the log_2_ fold-change of the relative abundance of each member within the community in response to each dietary compound compared to the base rich media (mGAM). The ratio of the relative abundance of *S. copri* over the sum relative abundance of the six Bacteroidaceae strains has been calculated for each condition. Data represents the average from three replicates. (**E**) Pie chart representing the proportion of dietary compounds from each category that significantly increase the relative abundance of *S. copri* in the community (FDR<0.05; Kruskal-Wallis with false discovery rate correction). (**F**) Total number of dietary compounds that decrease or increase the relative abundance of each commensal member within the community (n=3; FDR<0.05; Kruskal-Wallis with false discovery rate correction).

To determine which species in the community are capable of utilizing the different compounds, we tested each of the 21 individual community members for their capacity to grow on each of the 79 dietary glycans as a sole carbon source (n=1659 bacteria/glycan pairs; Extended Data Fig. 1c,d; Supplementary Table 2). We found that five Bacteroidales, *Bifidobacteria*, and *E. boltae* grew on most compounds (Extended Data Fig. 1c). Interestingly, despite *B. ovatus* growing on the largest number of compounds, and other Bacteroidaceae utilizing a similar number of compounds to *S. copri* (>25 % growth as a fraction of growth in mGAM), most glycans promoted *S. copri* growth in the community at the expense of the Bacteroidaceae (Fig. 1f, Extended Data Fig. 1c). Additionally, the capacity of the different species to utilize the glycans did not predict the direction of Prevotellaceae/Bacteroidaceae ratios within the community. For instance, pectic galactan is utilized by *S. copri*, *B. thetaiotaomicron, B. uniformis*, *B. ovatus*, and *B. caccae*, but when added, it leads to an increase in *S. copri* within the community and not the Bacteroidaceae species (Fig. 1d, Extended Data Fig. 1d). Further, *B. thetaiotaomicron*, *P. vulgatus*, and *S. copri* all utilize arabinan, when added, it leads to an expansion of *S. copri* and a depletion of the Bacteroidaceae members (Fig. 1d, Extended Data Fig. 1d). These experiments demonstrate that assessing the utilization capacity of dietary components by individual commensal species in monoculture does not reliably predict the overall community dynamics. Other factors of bacterial cross-talk, besides the complex carbohydrate utilization capacity, probably influence community structure and composition. These results demonstrate that complex carbohydrates and vitamins significantly impact gut community structure, especially the Prevotellaceae/Bacteroidaceae ratio.

### Community members influence Segatella copri and Bacteroidaceae competition

Our results indicate an inverse relationship within the community between *S. copri* and the Bacteroidaceae in the presence of most compounds. We next investigated whether *S. copri* and the Bacteroidaceae species compete within the community. We chose arabinan supplementation as a model to investigate these dynamics in detail, as arabinan strongly promotes *S. copri* over the Bacteroidaceae species, and the PUL specific for arabinan utilization is known for *S. copri*, *B. thetaiotaomicron*, and predicted for *P. vulgatus* ^20,24^. Moreover, *B. thetaiotaomicron*, *P. vulgatus*, and *S. copri* grow efficiently on arabinan (Extended Data Fig. 2a,b). Compared to mGAM, supplementation of arabinan led to a ∼35-fold increase in *S. copri* relative abundance (1.7 % (SD 0.25 %) to 59.6 % (SD 2.5%)) and >6-fold decrease in overall Bacteroidaceae (47 % (SD 2.4 %) to 7.3 % (SD 0.9%)), with *P. vulgatus* relative abundance decreasing to 1.4 % (SD 0.15%) (Fig. 2a). Thus, *S. copri* has a clear advantage within the community in the presence of arabinan (Fig. 2a,e). To determine if *S. copri* requires direct arabinan utilization to gain an advantage, we analyzed the gene expression of the arabinan-specific PUL14 of *S. copri* within the community by metatranscriptome analysis. The SusC/D-like gene pairs in the *S. copri* arabinan-specific PUL14 are upregulated up to ∼16-fold compared to mGAM alone, suggesting active arabinan metabolization by *S. copri* within the community (Fig. 2b, Supplementary Table 3). Of note, genes in the *B. thetaiotaomicron* arabinan PUL were also upregulated (Supplementary Table 3). Next, we tested whether direct utilization of arabinan is required for *S. copri* expansion within the community in the presence of arabinan. Deletion of SusC1 in the arabinan PUL14 renders *S. copri* unable to utilize arabinan ^20^ (Extended Data Fig. 2c). Indeed, unlike *S. copri* WT, the *S. copri ΔsusC1* mutant strain was unable to expand within the community in the presence of arabinan (Fig. 2c). These results show that *S. copri* gains a competitive advantage within the community by directly utilizing arabinan.

**Figure 2.**
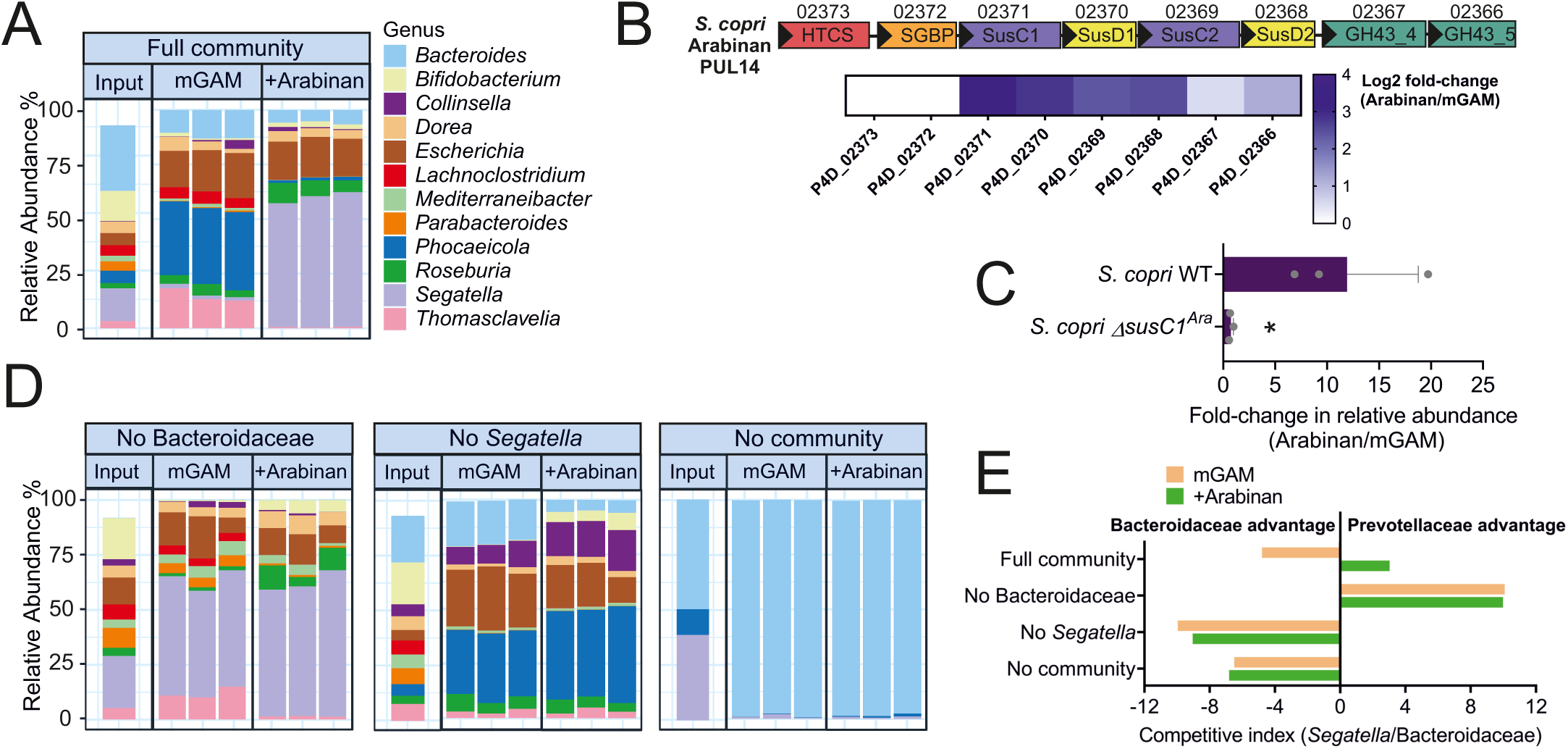
*S. copri* competitive advantage over Bacteroidaceae species in arabinan depends on other commensal members of the gut community. (**A**) Taxonomic bar plots depicting the relative abundance of the top 12 community genera at passage 3 and the input in mGAM. Data is from 16S rRNA amplicon sequencing with three biological replicates shown. (**B**) Upper panel: schematic representation of the genes in the *S. copri* arabinan utilization PUL14. Lower panel: heatmap displaying the log_2_ fold-change in gene expression of *S. copri* PUL14 genes at 20 h within the community in the presence of arabinan compared to mGAM alone. (**C**) Log_2_ fold-change of the relative abundance of *S. copri* WT and *ΔsusC1^Ara^* mutant within the community in the presence of arabinan compared to mGAM alone. Bars represent the average ±SD, *p-value<0.05 (Student t-test, n=3). (**D**) Taxonomic bar plots depicting the relative abundance of the top 12 community genera at passage 3 and the input, in the presence of arabinan or mGAM alone. Data is from 16S rRNA amplicon sequencing with three biological replicates shown. “No Bacteroidaceae” communities excluding six Bacteroidaceae species, “No *Segatella*” communities excluding *Segatella copri*, “No community” includes only six Bacteroidaceae species and *Segatella copri*. (**E**) Calculated competitive index between Prevotellaceae and Bacteroidaceae in community assemblies from A and D (See Methods). In A and D data is representative of n=2-7 independent experiments.

Next, to determine whether *S. copri* and Bacteroidaceae compete within the community, we assembled the communities by omitting either *S. copri* or all six Bacteroidaceae species. In line with a model of direct competition, we found that in the absence of Bacteroidaceae, *S. copri* occupies the full Bacteroidales niche both in mGAM and with arabinan (Fig. 2d,e). Conversely, Bacteroidaceae occupied the full Bacteroidales niche in both mGAM and with arabinan in the absence of *S. copri* (Fig. 2d,e). This indicates that *S. copri* and Bacteroidaceae species compete within the community and have a distinct advantage in the presence of different complex carbohydrates (e.g., arabinan vs starch). We hypothesized that *S. copri* could scavenge arabinan more efficiently than *P. vulgatus* and *B. thetaiotaomicron* to gain a competitive advantage within the community. To test this, we competed directly *S. copri* with the six Bacteroidaceae species in the absence of other community members. Surprisingly, *S. copri* was fully outcompeted by the Bacteroidaceae species, primarily *B. thetaiotaomicron* and *B. fragilis*, in the presence of arabinan when other commensals were absent (Fig. 2d,e; Extended Data Fig. 2d). Together, these results indicate that while direct arabinan utilization is necessary for *S. copri* expansion within the community, it is not sufficient; interactions with other community members are required for *S. copri* to dominate in the presence of a preferred complex glycan.

### S. copri expansion within the community depends on E. coli

We observed that *S. copri* was able to outcompete six Bacteroidaceae species within the complex 21-member community when arabinan was supplemented. However, in the absence of the broader community, *S. copri* was completely outcompeted by the same Bacteroidaceae, despite the presence of arabinan. To identify the community member(s) that provide a competitive advantage to *S. copri* in the presence of arabinan, we employed an add-in strategy where we competed *S. copri* with the six Bacteroidaceae species and added commensal members that displayed >0.1% relative abundance in the community in the presence of arabinan (Fig. 2a). Species from the remaining three phyla were tested; Bacillota (three Lachnospiraceae species or *Thomasclavelia*) or Actinomycetota (two *Bifidobacteria sp.* or *Collinsella*) did not alter the competitive advantage of Bacteroidaceae over *Segatella* in all conditions (Fig. 3a,b). Strikingly, addition of the Pseudomonadota member, *E. coli,* was sufficient to shift the competitive advantage in favor of *Segatella* in the presence of arabinan, increasing its relative abundance from 0.77 % (SD 0.4 %) in mGAM alone to 77.7 % (SD 2.3 %) in arabinan-supplemented mGAM (Fig. 3a,b). This shows that *E. coli* alone is sufficient to provide *S. copri* with a competitive advantage over Bacteroidaceae species in the presence of arabinan.

**Figure. 3.**
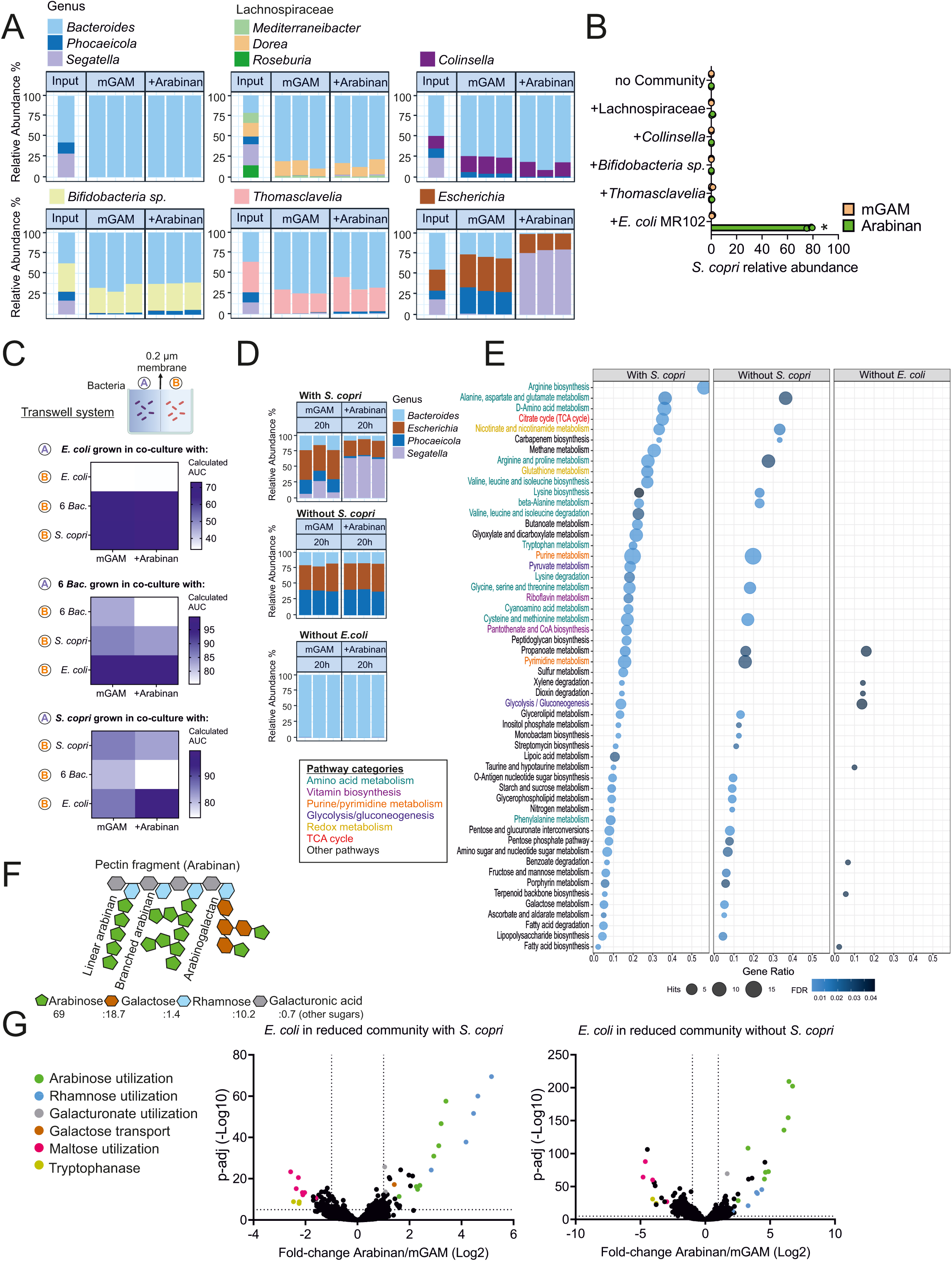
*E. coli* provides a competitive advantage to *S. copri* over Bacteroidaceae. (**A**) Taxonomic bar plots at the genus level showing the relative abundance of the community members in the input and at passage 3 with arabinan or mGAM alone. Data is from 16S rRNA amplicon sequencing with three biological replicates shown. (**B**) *S. copri* relative abundance in the competition with Bacteroidaceae, with the addition of different species in the presence or absence of arabinan. *p-value<0.0001 from t-test with correction for multiple comparisons, n=3. (**C**) Schematic of transwell co-culture platform (top). Area under the curve (AUC) calculated from growth curves in transwell co-cultures (bottom). Each species was co-cultured with itself as a monoculture control. The data is representative of two independent experiments. (**D**) Relative abundance of the genera in the different community assemblies: “with *S. copri*”, “without *S. copri*”, and “without *E. coli*”. Data is from 16S rRNA amplicon sequencing with three biological replicates shown. (**E**) Dot plot displaying enrichment of metabolic pathways upon arabinan supplementation compared to the mGAM control for each community assembly shown in (D). “Hits” indicates the number of significantly enriched metabolites within each pathway, while “Gene Ratio” denotes the proportion of significantly changed metabolites relative to the total metabolites in the pathway. Metabolic pathway enrichments with FDR < 0.05 were considered significant. n = 3. (**F**) Schematic representation of a pectin fragment depicting the monosaccharides constituting arabinan polysaccharides. (**G**) Volcano plots showing gene expression changes in *E. coli* MR102 within communities “with *S. copri*” or “without *S. copri*” in the presence of arabinan compared to mGAM control.

To determine the influence of *E. coli* on the growth of the Bacteroidales and vice versa, we performed trans-well co-culture assays allowing nutrient exchange (0.2 μm barrier) with different combinations of *S. copri*, *E. coli*, and the six Bacteroidaceae compared to monocultures (Fig. 3c, Extended Data Fig. 3). Interestingly, we observed that *E. coli* displayed improved growth when co-cultured with *S. copri* or the six Bacteroidaceae species in mGAM and in the presence of arabinan (Fig. 3c, Extended Data Fig. 3). Bacteroidaceae growth was modestly increased in the presence of *S. copri*, whereas co-culture with *E. coli* boosted Bacteroidaceae growth independent of media (Fig. 3c, Extended Data Fig. 3). *S. copri* growth was modestly inhibited by co-culture with Bacteroidaceae, and remarkably, *S. copri* growth was boosted when co-cultured with *E. coli* in the presence of arabinan compared to mGAM alone (Fig. 3c, Extended Data Fig. 3). These results are in agreement with *E. coli* and arabinan synergizing to promote *S. copri* predominance over Bacteroidaceae species.

### Metabolomics and metatranscriptomics of synthetic communities reveal cross-feeding between Bacteroidales and E. coli

To uncover potential nutrient exchange mechanisms, we investigated the contribution of the different families, Prevotellaceae (*S. copri*), Bacteroidaceae (six species), and Enterobacteriaceae (*E. coli*) to the metabolic and transcriptomic landscape of the communities. First, we performed targeted metabolomics on the different community assemblies with or without arabinan supplementation. We analyzed *S. copri* with the six Bacteroidaceae in the presence of *E. coli* (“with *S. copri*”), the six Bacteroidaceae in the presence of *E. coli* without *S. copri* (“without *S. copri*”), and *S. copri* with six Bacteroidaceae in the absence of *E. coli* (“without *E. coli*”) (Fig. 3d). Consistent with our previous results, only in the presence of *E. coli*, the community switches from Bacteroidaceae-dominated to *S. copri*-dominated when supplemented with arabinan, whereas the absence of *E. coli* or *S. copri* leads to a Bacteroidaceae-dominated community irrespective of arabinan supplementation (Fig. 3d). Consequently, we observed the largest number of significantly changed metabolic pathways, based on changes in the relative concentrations of associated metabolites, when arabinan is supplemented in the community that includes both *S. copri* and *E. coli* (“with *S. copri*”) (Fig. 3d,e: Supplementary Table 4). Metabolic pathways involved in amino acid metabolism, vitamin biosynthesis, the TCA cycle, glycolysis/gluconeogenesis, and redox metabolism were enriched in the community “with *S. copri*”, which suggests these metabolites are contributed by the expansion of *S. copri* (Fig. 3e). Fewer metabolic pathways were enriched in the community “without *S. copri*” in the presence of arabinan. Purine and pyrimidine metabolism, amino acid metabolism pathways, and nicotinate/nicotinamide metabolism were enriched to a similar degree as to that of the communities “with *S. copri*” but not the communities “without *E. coli*” suggesting these pathways are contributed by the presence of *E. coli* (Fig. 3e). The community “without *E. coli*” showed a minor metabolic response to arabinan, reflecting the minor changes in the Bacteroidaceae-dominated community (Fig. 3d,e). These results indicate that the shift from Bacteroidaceae to a *Segatella*-dominated configuration, modulated by arabinan and the presence of *E. coli*, is the primary factor influencing the metabolic landscape.

Next, harnessing the advantage of defined communities, we performed metatranscriptomics and captured the gene expression of the individual members specifically. We analyzed the transcriptomic signatures of the individual bacterial families within these community assemblies to gain deeper insight into the interactions between the different species. The analysis of *S. copri* gene expression in the presence of arabinan in the community “with *S. copri*” showed an upregulation of various nutrient acquisition genes including PULs, transporters, glycosyl hydrolases, tryptophan biosynthesis, vitamin biosynthesis, and genes of unknown function, and a downregulation of genes in PUL13, various amylase genes, and DNA-binding proteins, which is consistent with the metabolomics results and the shift from starch to arabinan utilization (Extended Data Fig. 4a; Supplementary Table 5). Notably, genes in the *S. copri* arabinan PUL14 were significantly upregulated at 8h in the presence of arabinan (between 1.8-1.9 fold-change; padj<0.0001) (Supplementary Table 5). Further, we analyzed the gene expression of the three most abundant Bacteroidaceae species in the community assemblies: *B. thetaiotaomicron*, *B. fragilis*, and *P. vulgatus*. Overall, the gene expression changes between the different Bacteroidaceae in the presence of arabinan in the different community assemblies were similar (Extended Data Fig. 4b-d; Supplementary Table 5). All three Bacteroidaceae differentially regulated nutrient acquisition genes, including PULs, transporters, and glycosyl hydrolases, while *B. fragilis* additionally upregulated fimbrial genes (Extended Data Fig. 4b-d; Supplementary Table 5). Interestingly, gene expression analysis of *E. coli* in communities with and without *S. copri* showed an upregulation of genes involved in arabinose, rhamnose, galacturonate, and galactose utilization, and a downregulation in genes involved in maltose utilization and tryptophanase in the presence of arabinan (Fig. 3f,g; Extended Data Fig. 4e; Supplementary Table 5). This indicates that *E. coli* is engaged in cross-feeding on monosaccharides deriving from polysaccharide degradation, and upon arabinan supplementation, it switches from maltose utilization (from starch) to arabinose, rhamnose, galacturonate, and galactose utilization (from arabinan) (Fig. 3f,g). Of note, we observed a downregulation of tryptophanase genes, including *tnaA,* responsible for indole synthesis in *E. coli* (Fig. 3f,g; Extended Data Fig. 4e; Supplementary Table 5). This is consistent with the report that *E. coli*’s indole production is inhibited through monosaccharide cross-feeding resulting from fiber degradation by *B. thetaiotaomicron* ^25^. These results indicate that *E. coli* cross-feeds on monosaccharides derived from arabinan degradation by *S. copri* in the “with *S. copri*” community, and by *B. thetaiotaomicron* and *P. vulgatus* “without *S. copri*”. Altogether, these results show that *E. coli* engages in beneficial cross-feeding from fiber degradation and leads to the expansion of *S. copri* in the presence of a preferred complex carbohydrate.

### Segatella sp. competitive advantage depends on preferred complex carbohydrates and Enterobacteriaceae species

We identified that various complex dietary glycans, in addition to arabinan, promote *S. copri* expansion in the community. We next evaluated whether complex glycans besides arabinan could give *S. copri* an *E. coli*-dependent competitive advantage. We chose plant-derived polysaccharides that promote *S. copri* within the community, as well as starch that promotes Bacteroidaceae (Fig. 1d). All eight additional plant-derived polysaccharides tested promoted *S. copri* over Bacteroidaceae in an *E. coli*-dependent manner, including the starch derivatives amylose and amylopectin, but not starch (Fig. 4a). This shows that *E. coli* synergizes with a variety of complex carbohydrates to provide *S. copri* with a competitive advantage over Bacteroidaceae. We next investigated whether this fiber-dependent interaction with *E. coli* is conserved beyond the *S. copri* HDD04 strain. Indeed, additional *S. copri* strains DSM18205^T^ and HDC01 (former *Prevotella copri* clade A) also gained an *E. coli*-dependent advantage in the presence of arabinan (Fig. 4b).

**Fig. 4.**
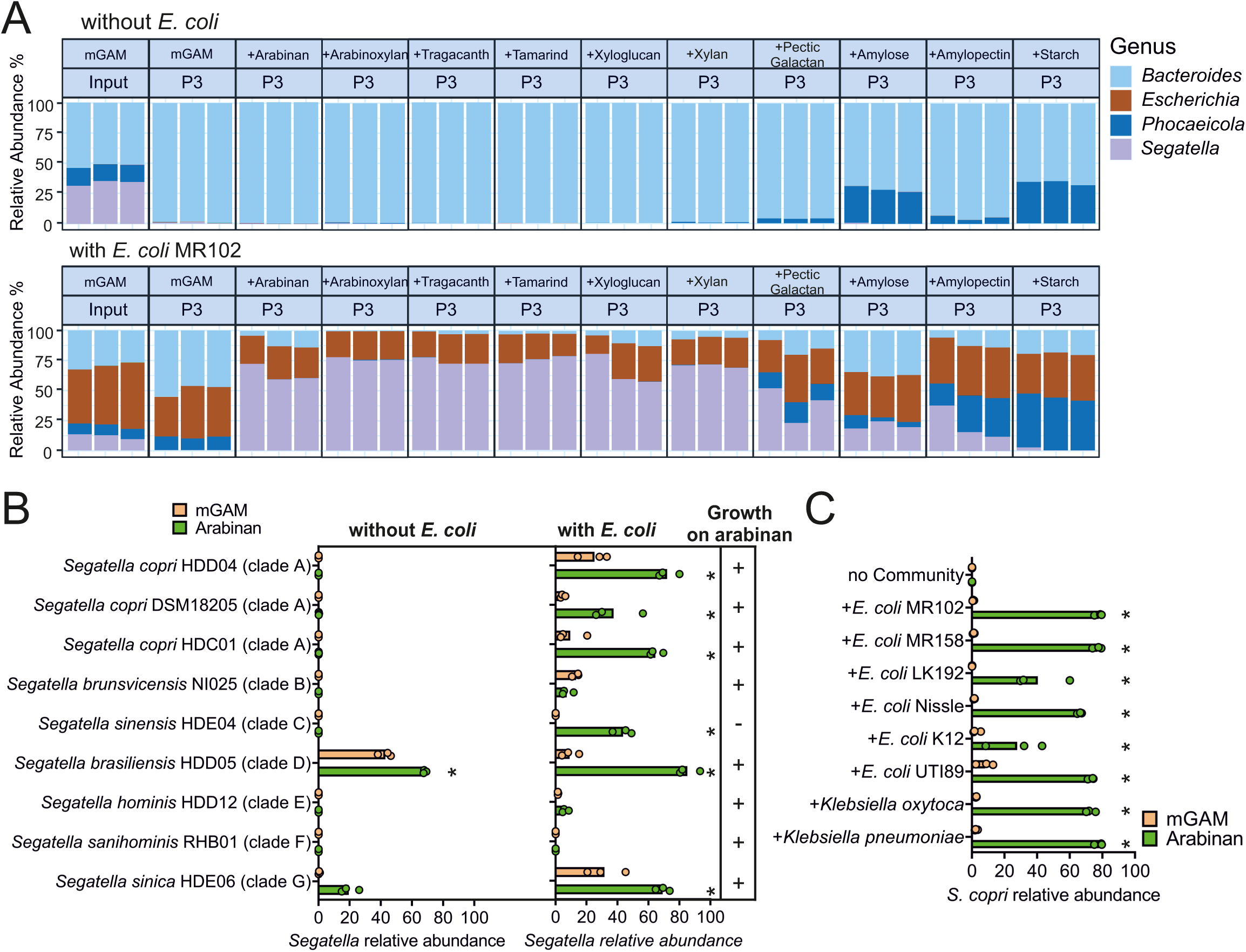
Enterobacteriaceae provide a competitive advantage to *Segatella sp.* in the presence of various complex polysaccharides. (**A**) Taxonomic bar plots at the genus level showing the relative abundance of the community members in mGAM or with a supplemented polysaccharide. Data is from 16S rRNA amplicon sequencing with three biological replicates shown. (**B**) Relative abundance of *Segatella sp.* in the competitions with Bacteroidaceae with or without *E. coli* MR102 in the presence or absence of arabinan. Growth on arabinan as a sole carbon source was tested for every *Segatella* strain in monocultures shown in the right panel. (**C**) *S. copri* relative abundance in the competitions with Bacteroidaceae, with the addition of different Enterobacteriaceae species in the presence or absence of arabinan. (B, C) *p-value<0.05 from t-test with correction for multiple comparisons, n=3.

Interestingly, the *E. coli*-dependent expansion can be observed beyond *S. copri* for other gut *Segatella* species such as *Segatella sinensis* HDE04 (former clade C) and *Segatella sinica* HDE06 (former clade G) showing that this interaction is conserved beyond *S. copri* (Fig. 4b; Extended Data Fig. 4f).

Finally, we tested whether this observed positive interaction between *S. copri* and *E. coli* occurs with genetically diverse *E. coli* strains. We found that *S. copri* relative abundance increased in the presence of arabinan with all tested *E. coli* strains, including other commensal isolates from healthy human donors (MR158 and LK192) ^26^, the probiotic strain *E. coli* Nissle, the lab strain *E. coli* K12, and the uropathogenic *E. coli* UTI89 (Fig. 4c). Strikingly, other Enterobacteriaceae species, namely a commensal isolate of *Klebsiella oxytoca* and a patient-derived *Klebsiella pneumoniae* strain, also lead to an increase in *S. copri* relative abundance in the presence of arabinan (Fig. 4c). Altogether, these results show that there is a broadly conserved mechanism of positive interaction between *Segatella sp.* and Enterobacteriaceae species in the presence of chemically diverse complex carbohydrates.

### Enterobacteriaceae species are prevalent in non-Westernized microbiomes

The dominance of *Segatella* species in non-Westernized gut microbiomes is strongly correlated with plant-based diets rich in fiber, but has remained causally unresolved ^13,27^. Our results show that Enterobacteriaceae, in addition to preferred complex carbohydrates, are an important component that promotes the shift towards a *Segatella*-dominated community. Therefore, we examined whether Enterobacteriaceae species are more prevalent in non-Westernized microbiomes, potentially supporting a predominance of Prevotellaceae. Specifically, we analyzed publically available gut metagenomes of ∼1000 healthy adult individuals with either Westernized or non-Westernized lifestyles (Fig. 5a; Supplementary Table 6). Consistent with previous reports, non-Westernized individuals harbor a higher prevalence, diversity, and abundance of Prevotellaceae compared to Westernized individuals (Fig. 5b; Extended Data Fig. 5a) ^15^. Interestingly, non-Westernized individuals also harbored a significantly higher prevalence, diversity, and abundance of Enterobacteriaceae species compared to Westernized individuals (Fig. 5c; Extended Data Fig. 5b). Finally, we observed a positive correlation between the relative abundance of Prevotellaceae and the number of Enterobacteriaceae species, and specifically, *S. copri* relative abundance positively correlated with the number of Enterobacteriaceae species in non-Westernized individuals (Extended Data Fig. 5c,d). Overall, non-Westernized microbiomes are richer in Enterobacteriaceae compared to Westernized microbiomes, and Prevotellaceae and Enterobacteriaceae are positively correlated in adults. Of note, the distinct Prevotellaceae-dominated and Bacteroidaceae-dominated microbiome signatures appear early in life (6-36 months of age), following an initial *Bifidobacteria*-dominated microbiome driven by exclusive milk consumption (0-6 months) ^28^. As reported, non-Westernized infants display a higher prevalence of Prevotellaceae starting from early life compared to Westernized infants (Extended Data Fig. 5e)^28^. Interestingly, we observed that Enterobacteriaceae prevalence is distinct in infants from different lifestyles. Despite a higher Enterobacteriaceae prevalence in the early life of Westernized infants, there is a significantly higher prevalence of Enterobacteriaceae in non-Westernized infants at 2-3 years of age (Extended Data Fig. 5e). This suggests that as Prevotellaceae becomes dominant, a prominent niche for Enterobacteriaceae also tends to form.

**Fig. 5.**
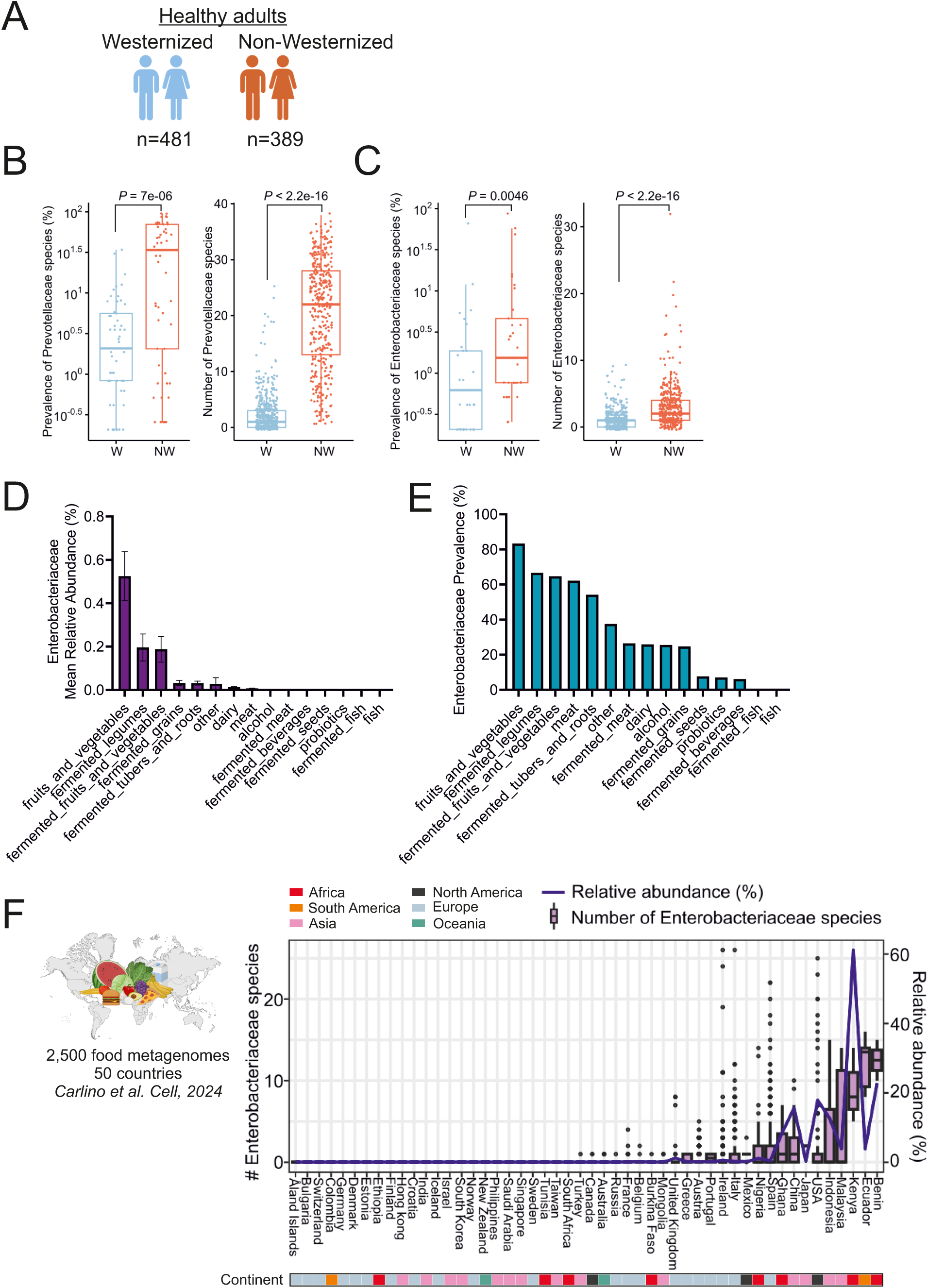
Enterobacteriaceae are prevalent in non-Westernized microbiomes. (**A**) Schematic representing the healthy adult cohort with Westernized and non-Westernized lifestyles (Supplementary Table 6). (**B**) Prevalence and number of Prevotellaceae species in the gut microbiome of Westernized (W) and non-Westernized (NW) healthy adults. P-values were determined using a two-tailed Wilcoxon rank-sum test. (**C**) Prevalence and number of Enterobacteriaceae species in the gut microbiome of Westernized (W) and non-Westernized (NW) healthy adults. P-values were determined using a two-tailed Wilcoxon rank-sum test. (**D, E**) Relative abundance (**D**) and prevalence (**E**) of Enterobacteriaceae in the different categories of food microbiomes. (**F**) Prevalence and relative abundance of Enterobacteriaceae species in food microbiomes from different countries, data from ^35^. Box plots were presented as: middle line, median; lower hinge, first quartile; upper hinge, third quartile; lower whisker, the smallest value at most 1.5× the interquartile range from the hinge; upper whisker, the largest value no further than 1.5× the interquartile range from the hinge; data beyond the whiskers are outlying points.

Enterobacteriaceae are generally a minor inhabitant of the healthy human gut in terms of abundance, but as facultative anaerobes, they are major fecal contaminants that can survive and spread through the environment, including foods ^29–34^. We wondered whether Enterobacteriaceae are more prevalent in the non-Westernized environment, specifically in food items. To address this hypothesis, we took advantage of an extensive dataset of food metagenomes from the curatedFoodMetagenomicData (cFMD) database ^35^. Strikingly, we found that raw fruits and vegetables, foods rich in plant fibers, are the foods with the highest relative abundance and prevalence in Enterobacteriaceae (Fig. 5d,e). We also observed that foods containing a significantly higher relative abundance and diversity of Enterobacteriaceae originate from countries characterized by predominantly non-Westernized lifestyles (Fig. 5f). Notably, the top five countries account for 69% of the relative abundance and 72% of the total number of Enterobacteriaceae species detected in food samples globally; these countries are also characterized by predominantly non-Westernized lifestyles (Fig. 5f). Altogether, these results show that Enterobacteriaceae are highly prevalent in non-Westernized microbiomes of healthy individuals and environmental settings such as foods. This points to food as a potential contributing factor in the distribution of Enterobacteriaceae, and is consistent with the experimental results demonstrating synergistic interactions between dietary components and Enterobacteriaceae that promote *Segatella* dominance.

## Discussion

The distinct prevalence of *Segatella sp.* in the gut microbiome of individuals with non-Westernized or non-industrialized lifestyles, as well as ancient human specimens, has been documented; however, the underlying mechanisms remain largely experimentally unexplored ^36^. This phenomenon is primarily attributable to the historical under-sampling of these populations, coupled with the fastidious nature of *Segatella sp.* regarding culturing, genetic manipulation, and establishment within experimental models. Nevertheless, the predominance of *Segatella sp.* in global populations emphasizes the necessity for a more comprehensive understanding of these bacteria. Despite this, the reasons underlying the preference for Bacteroidaceae over *Segatella* in Westernized microbiomes remain ambiguous. While a correlation with dietary patterns has been suggested ^27,37^, other studies have failed to find a link between *Segatella* abundance and short-term dietary interventions or habitual diets among cohorts from the same geographical region, indicating that diet alone does not explain *Segatella* predominance ^13,16,18,38^. Our synthetic gut community design, which facilitates colonization by *S. copri*, coupled with our previously established genetic tools and *Segatella* isolates, has illuminated the mechanisms governing *Segatella* predominance in the gut microbiota. Utilizing our experimental model, we demonstrate that *Segatella* preferentially assimilate a wide variety of complex dietary glycans, which act synergistically with Enterobacteriaceae to promote its expansion within a gut microbial community.

A competitive relationship between *Segatella* and *Bacteroides* species has been suggested; however, it has not been experimentally validated ^39^. Previous experimental setups failed to consistently demonstrate *S. copri* predominance over *Bacteroides* species in gut communities. In reported studies involving either 1:1 competitions or within commensal communities, *Bacteroides sp.* generally outcompete *S. copri* or both species co-exist ^40–43^. Whereas microbiota accessible carbohydrates (MACs) facilitate *S. copri* colonization, they do not confer a competitive advantage to *S. copri* over *B. thetaiotaomicron* alone ^42^. We observed that within the commensal community, *S. copri* and Bacteroidaceae engage in competition, with their dynamics influenced by conditions that favor one organism over the other. Notably, Bacteroidaceae prevailed over *S. copri* in the presence of various complex glycans, but *S. copri* is able to overcome this competitive disadvantage by leveraging support from other community members. The introduction of the community, or solely *E. coli* provided a single *Segatella* strain with a competitive advantage over multiple Bacteroidaceae species (including *Bacteroides* and *Phocaeicola*) in the presence of a preferred polysaccharide. Although, nutrient-mediated mutualistic or antagonistic pairwaise interbacterial relationships within the microbiome have been well documented ^44–46^, this study provides an example of more complex interactions, in which both nutrient exchange and bacterial crosstalk are needed for *Segatella* to dominate the microbial community.

The utilization of complex dietary glycans has been extensively explored in *Bacteroides* species, yet significantly less is known for *Segatella*. Our findings indicate that besides its specific preference for plant-derived fibers, *S. copri* also exhibited expansion in response to most fibers derived from microbial and algal sources. As previously noted, only a small portion of animal or host-derived glycans enhanced the abundance of *S. copri* in the community ^21,42^. This suggests that *S. copri* possesses a broad preference for complex glycans from various sources. Interestingly, *S. copri* showed a complex glycan utilization ability comparable to other Bacteroidaceae, despite having a significantly smaller number of glycan processing enzymes (29 predicted PULs and ∼40 CAZymes for *S. copri* HDD04) when compared to *B. thetaiotaomicron* and *B. ovatus*, which have around 100 PULs and hundreds of CAZymes ^20,21,47^. This may imply a greater versatility of *S. copri*’s PULs and CAZymes or indicate the existence of additional unknown loci involved in diverse complex glycan utilization. Indeed, genes encoding for polysaccharide catabolism enzymes have also been found outside of PULs in *Bacteroides* ^46^. Likewise, predictions regarding growth on certain polysaccharides for *Segatella* strains demonstrated that these strains could grow on specific polysaccharides even in the absence of predicted dedicated PULs ^21^. This provides an important resource for future studies on *Segatella* carbohydrate utilization and its effects on colonization. Besides dietary glycans, vitamin B3 and B5 promote *S. copri* expansion while decreasing Bacteroidaceae abundance within the community (Fig. 1). *S. copri* and various Bacteroidaceae species are predicted to synthesize most B vitamins, which are essential for their colonization ^48,49^. It has been shown that dietary supplementation with these vitamins increases *Prevotella* and decreases *Bacteroides* relative abundance in lactating women ^50^. This implies that these bacteria may compete for dietary vitamins in addition to glycans. However, the mechanisms underlying competition for these vitamins, particularly from dietary intake, have yet to be fully understood.

Studies have shown that *Bacteroides sp.* engage in mutualistic cross-feeding with pathogenic and commensal Enterobacteriaceae species ^25,51–53^. Enterobacteriaceae are generally unable to degrade complex carbohydrates and rely on fiber-degraders to acquire simple sugars in the gut. Our results show that similarly *S. copri* engages in mutualistic cross-feeding with *E. coli*.

We show that *E. coli* synergizes specifically with arabinan to provide *S. copri* with a competitive advantage over Bacteroidaceae. Interestingly, this mutualistic interaction with *S. copri* is conserved among Enterobacteriaceae and synergizes with various types of polysaccharides. Therefore, the presence and diversity of Enterobacteriaceae species within the microbiome could play an important role in determining *Segatella* establishment. Indeed, our human stool metagenomics analysis shows that Enterobacteriaceae prevalence, diversity, and abundance in non-Westernized microbiomes are significantly higher than in Westernized microbiomes. The richness of Enterobacteriaceae in non-Westernized microbiomes could be generally a consequence of the interaction with the environment ^30^. This could be potentially explained by a larger exchange of these strains with the environment in non-Westernized populations due to their considerable presence in the environment ^54^. Our analysis of an existing global dataset provided insights into the distribution of these bacteria across various food types in different countries ^35^. We observed a higher presence of Enterobacteriaceae in several countries with reported non-Westernized lifestyles, and specifically in raw fruits and vegetables. This is consistent with the correlation of *Segatella* prevalence with plant-based diets and anti-correlation with processed foods in westernized populations. Additionally, Enterobacteriaceae strain exchange can occur with other environmental sources, such as drinking water ^55^. In fact, extensive Enterobacteriaceae strain exchange between individuals has been associated with contaminated drinking water in urban informal settlements in Nairobi, Kenya ^33^. Finally, our findings suggest that establishing microbiota signatures by dominant gut Bacteroidales goes beyond simple dietary ingredients. It rather involves nutrient-driven commensal crosstalk and competition, and through environmental exposures that may promote microbiome enrichment.

## Supplementary materials

Extended Data Figure 1.

Extended Data Figure 2.

Extended Data Figure 3.

Extended Data Figure 4.

Extended Data Figure 5.

Supplementary Table 1. Community strains

Supplementary Table 2. Dietary components

Supplementary Table 3. Metatranscriptomics at 20h related to Figure 2

Supplementary Table 4. Metabolomics data related to Figure 3

Supplementary Table 5. Metatranscriptomics at 8h related to Figure 3

Supplementary Table 6. Human metagenomic data used for analysis in Figure 5

## Materials and Methods

### Bacterial strains and culture conditions

All strains used in this study are listed in Supplementary Table 1.

Gut commensal isolates and community assemblies were cultured in a flexible anaerobic chamber (Coy Laboratory Products) containing 10% CO2, 5% H2, and 85% N2 at 37°C in liquid modified Gifu Anaerobic broth (mGAM, Hyserve #05433) and grown on brain heart infusion (BHI; Oxoid #CM1135B) agar (BACTO^TM^ agar, BD # 214010) supplemented with 10% defibrinated sheep blood (Thermo Scientific, # 10516583) and 1μg/ml Menadione (vitamin K_3_, Sigma-Aldrich # M5625).

### Defined gut community design and assembly

To select members of the commensal community, isolates were chosen based on several criteria: they are commonly reported as human gut isolates and generally recognized as commensals; they grow well (within 24 h) in mGAM broth under anaerobic conditions, and they remain stable within the community after the first passage. Furthermore, the phylum-level composition of the community reflects that of the human gut, with proportional representation including representatives from eight major human gut bacterial families and with a species richness comparable to that of existing model communities. The final community is based on several iterations informed by previous studies and on the successful establishment of *Segatella sp.* ^11,41,56–58^.

To assemble communities, strains were grown overnight in mGAM from single colonies. Each strain was adjusted to a final OD_600_ of 0.01 and pooled into fresh media. After 24 hours of growth (P0), the strains were passaged at a 1:100 dilution into fresh mGAM or mGAM supplemented with the corresponding dietary component. Communities were harvested at passage 3 (P3), identified as the time-point when the community composition reached stability. At P3, 1 ml of the culture was centrifuged to pellet the cells, which were then immediately frozen and stored for subsequent analysis, unless otherwise specified.

### Screen of dietary components

The stock solutions of the dietary components were prepared as indicated in Table S2.

To prepare the supplemented media, 2x mGAM, dissolved in MilliQ water and filter sterilized, was pooled with the stock solution of the dietary compound (v/v) (Supplementary Table 2). Dietary glycans were prepared as described and used at final concentrations ranging from 0.04% to 0.2%, adjusted based on properties such as gelling, solubility, viscosity, and accessible amounts. Vitamins were prepared separately as indicated and used at concentrations corresponding to those previously tested *in vivo* ^59^. The media was aliquoted into sterile 96-deep well plates and inoculated with the community assembly previously grown for 24 hours in mGAM (P0); cultures were adjusted to a starting OD_600_ of 0.01. The plates were covered with breathable membranes and passaged into fresh media until passage 3 using an Eppendorf epMotion® 96 with mixing. The endpoint OD_600_ at each passage was measured for all conditions and normalized to media-only background.

### Flow cytometry analysis

For flow cytometry analysis, 500 μl of community culture was collected at P3 in microcentrifuge tubes and washed twice with 1 mL PBS (4,500 rpm, 4 min at 4°C). After the second wash, cells were re-suspended in 1 ml propidium iodide/DAPI 1µg/ml solution, 25μl of cell counting control beads were added, and the cells were gated based on forward and side scatter to exclude debris. Viability gating was performed using DAPI-positive cells to identify total bacteria and PI staining to distinguish dead or membrane-compromised cells (PI-positive). Absolute cell counts were calculated by comparing the number of bacterial events to the known concentration and number of counting beads detected, according to the manufacturer’s instructions.

### Growth on sole carbon source

The stock solution for growth on complex dietary glycans as sole carbon source were prepared as indicated in Table S2. A 2x concentration of filter sterilized minimal media was prepared as in ^20^ and pooled with each dietary glycan stock solution (v/v). The commensal strains were grown overnight in mGAM from single colonies and inoculated into the minimal media at a starting OD_600_ of 0.05 in 96-well plates. The cultures were grown for 70 hours at 37°C with OD_600_ measurement at 1 hour intervals after 45 s of orbital mixing using a Biotech Epoch2 reader.

To calculate the percent growth, the OD_max_ at each condition was calculated as a fraction of the OD_max_ for each strain in mGAM:

Percent growth = (ODmax minimal media/ODmax mGAM) x 100.

### Transwell co-culture experiments

Strains were grown overnight in mGAM from a single colony, and were used to inoculate 800µl of fresh mGAM or mGAM supplemented with arabinan at a starting OD_600_ of 0.05. The six Bacteroidaceae strains were pooled to a final starting OD_600_ of 0.05. Co-culture experiments were conducted using the Cerillo Duet Co-Culture Platform, which enables real-time monitoring of bacterial interactions while maintaining fluidic contact between compartments through a porous physical barrier (0.2 µm size exclusion filter). The Duet devices were inoculated with the respective bacterial strains designated for co-culture (800 µL volume per well). The plates were incubated for 48 h at 37 °C with OD_600_ measurement at 1 hour intervals after 45 s of orbital mixing using a custom Cerillo plate template on a Biotech Epoch2 reader. Area Under the Curve (AUC) for each growth curve was calculated using GraphPad Prism 8 with a Y=0 baseline for all curves.

### Genomic DNA extraction and amplicon 16S rRNA sequencing

Genomic DNA extraction from community samples was performed using the ZymoBIOMICS™ DNA Miniprep Kit, following the manufacturer’s protocol. For 16S rRNA amplicon sequencing, the V4 hypervariable region of the 16S rRNA gene was amplified using primers F515 (GTGCCAGCMGCCGCGGTAA) and R806 (GGACTACHVGGGTWTCTAAT), based on the established protocol ^60^. Briefly, input DNA was normalized to 25 ng/μl prior to PCR amplification, with unique 12-base Golay barcodes incorporated via specific primers (Sigma) to enable sample multiplexing.

Following PCR, amplicons were pooled and normalized to 10 nM before sequencing on an Illumina MiSeq platform using 250 bp paired-end reads (PE250). Raw sequencing reads were demultiplexed using idmp (https://github.com/yhmu/idemp) according to barcode sequences. Library processing included merging paired-end reads, quality filtering, dereplication, singleton removal, denoising, and chimera checking using the USEARCH pipeline version 11.0.66751.

Specifically, reads were merged with the fastq_mergepairs command (parameters: *maxdiffs=30, pctid=70, minmergelen=200, maxmergelen=400*), followed by quality filtering with *fastq_filter* (*maxee=1*). Unique sequences were identified using *fastx_uniques* with a minimum unique size of 2. Biological sequences (ASVs/zOTUs) were predicted and chimeras removed using *unoise3* (parameters: *minsize=10, alpha=2*). Amplicon quantification was performed with *usearch_global* (*strand=plus, id=0.97, maxaccepts=10, top_hit_only, maxrejects=250*).

Taxonomic assignment was carried out using *Constax2*, which integrates classifiers RDP, SINTAX, and BLAST, referencing the GreenGenes database. Resulting taxonomic data were summarized into a biom file for visualization in phyloseq and further downstream analyses. Operational taxonomic units (OTUs) with an abundance below 0.5% across all samples were excluded. A closed-reference database comprising the defined community members was employed for classification.

### Competitive index calculation

The competitive index between Prevotellaceae (*Segatella copri*) and Bacteroidaceae (six species) was calculated using their relative abundances derived from 16S amplicon sequencing data, according to the following equation:

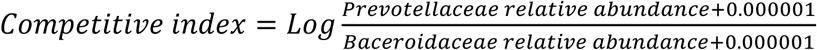

### qPCR quantification

The relative abundance of all bacteria was monitored at indicated time-points by qPCR of purified DNA using species-specific primers: for *B. thetaiotaomicron* forward (5’ GAGGGTGTCGTATTTCCGAAGG 3’) and reverse (GTTCCCTGATCCAGTGTGTTGG), for *B. caccae* forward (5’ GTGCAGAGTGGCATACTGAAAG 3’) and reverse (5’ AGTGTAGGCTTTCGTACCGTTC 3’), *B. ovatus* forward (5’ TTCCCGGAGAGTAAGCTAACAG 3’) and reverse (5’ GGAACAGTACAAGACCCTTTGG 3’), *B. uniformis* forward (5’ GCTCATACTGAAATGGGAGGAC 3’) and reverse (5’ GGTCAGGTAGGCAATGAGAATC 3’), *P. vulgatus* forward (5’ GAGTCGCGGACTGCATTATAC) and reverse (5’ ACACCCTCTTGGTGGATTCTG 3’), and *B. fragilis* forward (5’ AGAAGTATGGCATGGTCTTCTGG 3’) and reverse (5’ TAAGGCTGCGTGCAAATCAG 3’). qPCR was performed using a CFX96 instrument (BioRad) and SYBR FAST universal master mix (KAPA Biosystems). Mean strain quantities were calculated using a standard curve obtained from using the primers against purified genomic DNA from each specific strain, and relative changes were calculated with the efficiency-corrected ΔCq method ^61^.

### RNA processing and metatranscriptomics sequencing

RNA was processed and sequencing performed as in ^44^. Briefly, communities were collected after 8h and 20h of growth after passaging in mGAM and mGAM+arabinan, a biomass of 4 units of OD_600nm_ was collected 5:1 (v/v) with stop solution (95 % Ethanol, 5 % phenol) and snap frozen in liquid nitrogen. Bacterial total RNA was extracted by hot phenol method followed by a DNase treatment. Quality of the RNA was determined by bioanalyzer at genome analytics core facility of the Helmholtz Centre for Infection Research (HZI).

For metatranscriptomics sequencing, three biological replicates per condition were processed. Total RNA was subjected to ribosomal RNA depletion by Ribo-Zero Plus Microbiome rRNA Depletion Kit (Illumina) and cDNA libraries were generated by using Illumina Stranded Total RNA Prep (Illumina) following manufacturer’s instructions.

Generated libraries were sequenced at ∼100-200 million reads sequencing depth paired-end 2×150bp. To obtain species specific mapped reads, reads were aligned to the genomes of the specific members in community assemblies by the use of READemption (2.0.0) and *--crossalign_cleaning* command to remove reads mapped to multiple species ^62^. For each species, differentially expressed genes between mGAM and mGAM+arabinan conditions were calculated using DESeq2 (https://pubmed.ncbi.nlm.nih.gov/25516281/) that utilizes Wald test to determine the P-value and the Benjamini-Hochberg to correct p-values for multiple testing (P-adj) (https://pubmed.ncbi.nlm.nih.gov/25516281/).

## Targeted metabolomics of communities

### Targeted Metabolomics: Isotope ratio approach via LC-MSMS

For metabolomics analysis, the isotope ratio approach via LC-MSMS was used ^63^. Briefly, communities were collected after 20h of growth after passaging in mGAM and mGAM+arabinan. A biomass of OD_600_ of 1 for each community was collected and diluted quickly in chilled PBS to OD_600_ of 0.5. For targeted metabolomics, culture aliquots were vacuum-filtered on a 0.45-μm pore size filter (Merck Millipore #HVLP02500). Filters were immediately transferred into a 40:40:20 (v-%) acetonitrile (Honeywell # 14261-1 l)/methanol (VWR # 83638.320)/water extraction solution at −20 °C. Filters were incubated in the extraction solution for at least 30 min at −20 °C. Subsequently, metabolite extracts were centrifuged for 15 min at 13,000 RPM at −9 °C and the supernatants were stored at −80 °C until analysis.

Targeted metabolite relative concentrations were obtained from the ratio of the 12C signal of the sample and 13C signal of the internal standard (Supplementary Table 4). For metabolomics data analysis, the relative concentrations data were log10 transformed for further statistical analysis and converted to z-scores for multivariate analyses of the total metabolomic data. Metabolite enrichment analysis were performed using MetaboAnalyst 6.0 (https://www.metaboanalyst.ca). For pathway enrichment analysis, KEGG IDs and the relative concentrations of metabolites were utilized to identify enriched metabolic pathways based on the *Escherichia coli* K-12 MG1655 KEGG pathway database. The number of Hits, indicating the count of enriched metabolites per pathway, and the Gene Ratio, defined as the proportion of enriched metabolites relative to the total metabolites in each pathway, were calculated. Statistical significance was assessed using multiple t-tests, followed by false discovery rate correction.

### Metagenomics data analysis

To assess the global prevalence, richness, and abundance of Prevotellaceae and Enterobacteriaceae in human populations, we analyzed 870 publicly available shotgun metagenomes (Supplementary Table 6) from nine countries representing diverse lifestyles. Using MetaPhlAn 4 with default settings ^64^, we profiled microbial species and estimated relative abundances, retaining only species with ≥0.0001 abundance to reduce low-abundance noise. All samples were selected from healthy adult males and females (16 – 65 years), defined as free from self-reported diseases or medical interventions, to represent general populations. Each metagenome was classified as originating from either a Westernized or non-Westernized population (Supplementary Table 6). Briefly, the term “non-Westernized” describes a population practicing a traditional lifestyle relating to factors such as diet, hygiene, and medical healthcare. Whereas, Westernization is a process of incremental urbanization transitioning to controlled food production chain, increased hygiene, and accessibility to modern medicals. The concept of “Westernization” and “non-Westernization” was adopted here to demarcate populations by combining at least the majority of the factors discussed above, as described previously ^65,66^.

To identify Enterobacteriaceae members in food samples, we downloaded 2,483 taxonomic profiles from the most comprehensive food microbiome database – curatedFoodMetagenomicData (cFMD) (https://github.com/SegataLab/cFMD), and only retained taxa belonging to Enterobacteriaceae. Briefly, cFMD integrates 1,950 newly sequenced and 583 public food metagenomes from 50 countries, covering 15 food categories which can be further grouped into 107 types or 358 subtypes ^35^. The prevalence and relative abundance of Enterobacteriaceae members were examined on the basis of countries and continents. The relative abundance values of each Enterobacteriaceae members were obtained directly from the taxonomic profiles.

## Statistics and analysis

Statistical analysis was performed using GraphPad Prism 8 and R Software R 4.5.0 (www.r-project.org). Details of statistical tests used, sample size indicated as “n”, definition of means, error bars, and significance are provided in figure legends and in method details.

## Author contributions

T.S., Y.E.M., and C.T. conceived and designed experiments. C.T., Y.E.M, A.G., and L.O. performed experiments. J.R. and H. L. processed metabolomics data and analyzed raw mass data.

K.D.H. and N. S. analyzed metagenomics data and provided essential data and analysis tools. A.T. provided essential support, infrastructure, reagents, and analysis tools. T.S., Y.E.M., and C.T. wrote the paper. All authors read and edited the paper.

## Inclusion and ethics

We are committed to promoting diversity, equity, and inclusion in all aspects of our research to ensure fair representation and unbiased outcomes.

## Acknowledgements

We would like to thank Lisa Maier (Tübingen University) for the helpful discussions on community assembly. We thank Alexandra Koumoutsi, Carlos Voogdt, Diënty Hazenbrink, and Samir Giri (EMBL, Heidelberg) for helpful discussions and support. We thank Francesco Asnicar (University of Trento) for discussions on large dataset metagenomics analysis, and Pavaret Sivapornnukul and Xiayu Wang for assistance with plot generation. We thank Agata Bielecka, Marie Wende, and Till Robin Lesker for assistance with the preparation of RNA sequencing libraries, sampling, and 16S rRNA analysis. C.T. was awarded an HZI-EMBL Postdoctoral grant. TS was supported by the Deutsche Forschungsgemeinschaft (DFG, STR 1343/15 and EXC 2155 – project number 390874280).

